# What limits local ancestry inference at low divergence: a feasibility threshold, a metric that conceals failure, and a deficit of input more than architecture

**DOI:** 10.64898/2026.07.30.741148

**Authors:** Qing Tian

## Abstract

Local ancestry inference assigns each position along an admixed chromosome to a source population, underpinning admixture mapping, ancestry-specific association testing and admixture dating. Validation is almost exclusively on continentally divergent sources (Hudson’s *F_ST_≈* 0.1) and coalescent simulations; we examine both restrictions. Across *F_ST_* from 0.0022 to 0.243 we compare five methods — two likelihood baselines, RFMix, FLARE and a dilated convolutional network — on identical sites with exact ground truth, and on 11 real 1000 Genomes pairs. Three findings follow. First, a feasibility floor: at *F_ST_*= 0.0022 no method exceeds 0.575, and CHB/CHS at *F_ST_*= 0.00042 yields at best 0.551. Pairs motivating fine-scale analysis, such as northern versus southern Han, fall below it. Second, per-site accuracy conceals a failure of tract structure: the most accurate method per site produces 78.8*×* too many tracts, implying an admixture time 61.2*×* too old, which Viterbi decoding removes at no cost to accuracy (+0.0002). Third, the simulated lead does not survive real data, and the deficit is one of input more than of the architectures we varied: attention, state-space layers, capacity, objective and self-supervised pretraining each move accuracy by at most 0.006, while supplying the haplotype information the released tools receive recovers +0.031 on 8 of 8 pairs below *F_ST_* = 0.04 and nothing above it — necessary but not sufficient, since the network still trails on 10 of 11 pairs. Two quantities usually held fixed matter more than architecture: the statistic summarising reference matching, and reference panel size, which no method is near saturating.

## Introduction

An individual descended from a recent admixture event carries a chromosome that is a mosaic of segments, each inherited from one of the contributing source populations. Local ancestry inference (LAI) is the problem of recovering that mosaic: assigning to every position in the genome the source population from which it descends. LAI is a prerequisite for admixture mapping, for ancestry-specific effect-size estimation in genome-wide association studies of admixed cohorts, for controlling population structure at fine scales, and for dating admixture events from the length distribution of ancestry tracts.

The methods in routine use fall into two families. The first models the mosaic explicitly as a hidden Markov process along the chromosome, with emissions given by the allele frequencies or haplotype composition of reference panels; HAPMIX (Price et al. 2009), RFMix (Maples et al. 2013), Loter (Dias-Alves et al. 2018) and FLARE (Browning et al. 2023) are representative. The second family learns the mapping directly from simulated training data, treating LAI as a sequence-labelling problem; recent examples adapt convolutional segmentation architectures from computer vision (Montserrat et al. 2020), and exchangeable architectures have been applied more broadly to population-genetic inference (Chan et al. 2018).

Both families have been developed and validated in a narrow region of parameter space. The canonical benchmarks are admixture between African, European and Native American sources, as in African-American and Latino cohorts. These source pairs are separated by Hudson’s *F_ST_* of roughly 0.1 to 0.15. In that regime the per-site information available to distinguish sources is substantial, and published accuracies exceed 0.95.

Many admixture events of scientific interest do not resemble this. Population structure within East Asia is dominated by a north–south cline among Han Chinese. Measured on chromosome 22 of the 1000 Genomes phased release (Byrska-Bishop et al. 2022), Han Chinese in Beijing and Southern Han Chinese differ at *F_ST_* = 0.00042, some 260-fold below the European/East Asian comparison of 0.10924 computed on the same sites. Within-Europe pairs are comparable or smaller: Iberian versus Tuscan at 0.00152, British versus Utah-European at 0.00003. Divergences an order of magnitude larger correspond not to within-Han structure but to Han versus neighbouring East Asian groups: Kinh at 0.00650, Japanese at 0.00826, Dai at 0.00826. Whether the LAI machinery transfers to this regime is an open question of direct practical consequence: if it does not, published inferences of fine-scale ancestry tracts within such populations rest on an untested assumption.

This gap has begun to be addressed. Medina Tretmanis et al. (2026) recently evaluated existing and novel neural LAI methods under deliberately difficult conditions, including downsampled reference panels, intracontinental admixture, and temporally distant admixture events, and reported that inference power is comparatively low for intracontinental scenarios. Their study establishes that the problem is hard in this regime and compares architectures across a set of named scenarios.

Several questions remain open, and they are the subject of this work. Divergence is treated categorically (intercontinental versus intracontinental) rather than as a continuous axis, so a practitioner cannot look up the *F_ST_* of their own source pair and read off an expected accuracy, still less determine whether inference is feasible at all. Nor is it known whether the advantage of a learned method over classical smoothing is uniform across divergence or concentrated within it. Beyond those, two questions concern the evaluation itself rather than the methods: whether per-site accuracy, the metric these comparisons almost always report, tracks the quantities downstream analyses actually consume; and whether a comparison conducted in simulation predicts the same comparison on real haplotypes. We find that it does not, in both cases. That failure then raises a fourth question, which turns out to have the most practical answer: whether the learned method’s shortfall on real haplotypes reflects learned inference itself, or merely the representation it was given. Released tools consume reference haplotypes, whereas a frequency-only network does not. We find the shortfall is representational, that closing it is necessary but not sufficient, and that two quantities usually held fixed in such comparisons, the statistic used to summarise reference matching and the size of the reference panel, account for more of the difference between methods than any architectural choice we tested.

We address these questions as follows. We simulate two-way admixture across a gradient of source divergence spanning 0.0022 to 0.243, using a mosaicking construction in which local ancestry labels are exact rather than inferred. On identical evaluation windows we compare five methods: a windowed likelihood classifier, the same emissions smoothed by a hidden Markov model, the released implementations RFMix and FLARE, and a dilated convolutional network trained for per-site segmentation. We then repeat the comparison on real 1000 Genomes haplotypes, score the same predictions by tract structure rather than per-site accuracy, and measure sensitivity to phase switch error and to divergence misspecification. Finally we give that network the class of information the released tools receive (local haplotype-matching summaries against each reference panel) and repeat the real-data comparison. That identifies the divergence range within which the missing information matters, and two further comparisons bound its interpretation: summarising reference matching by contiguity rather than by agreement rate, and varying the reference panel size that every other experiment holds at a single value. Two configurations of the network are compared throughout, and we name them consistently from here on: the frequency-only network, which receives allele frequencies and their likelihood ratio, and the haplotype-aware network, the same architecture with four haplotype-matching channels appended. “The network” unqualified means the frequency-only configuration. Fig. 1 previews the argument panel by panel: the construction that makes the labels exact (A), the constraint divergence places on it (B), the discrepancy between per-site accuracy and tract structure (C), and the contrast between what changing the architecture achieved and what changing the input achieved (D).

**Figure 1.**
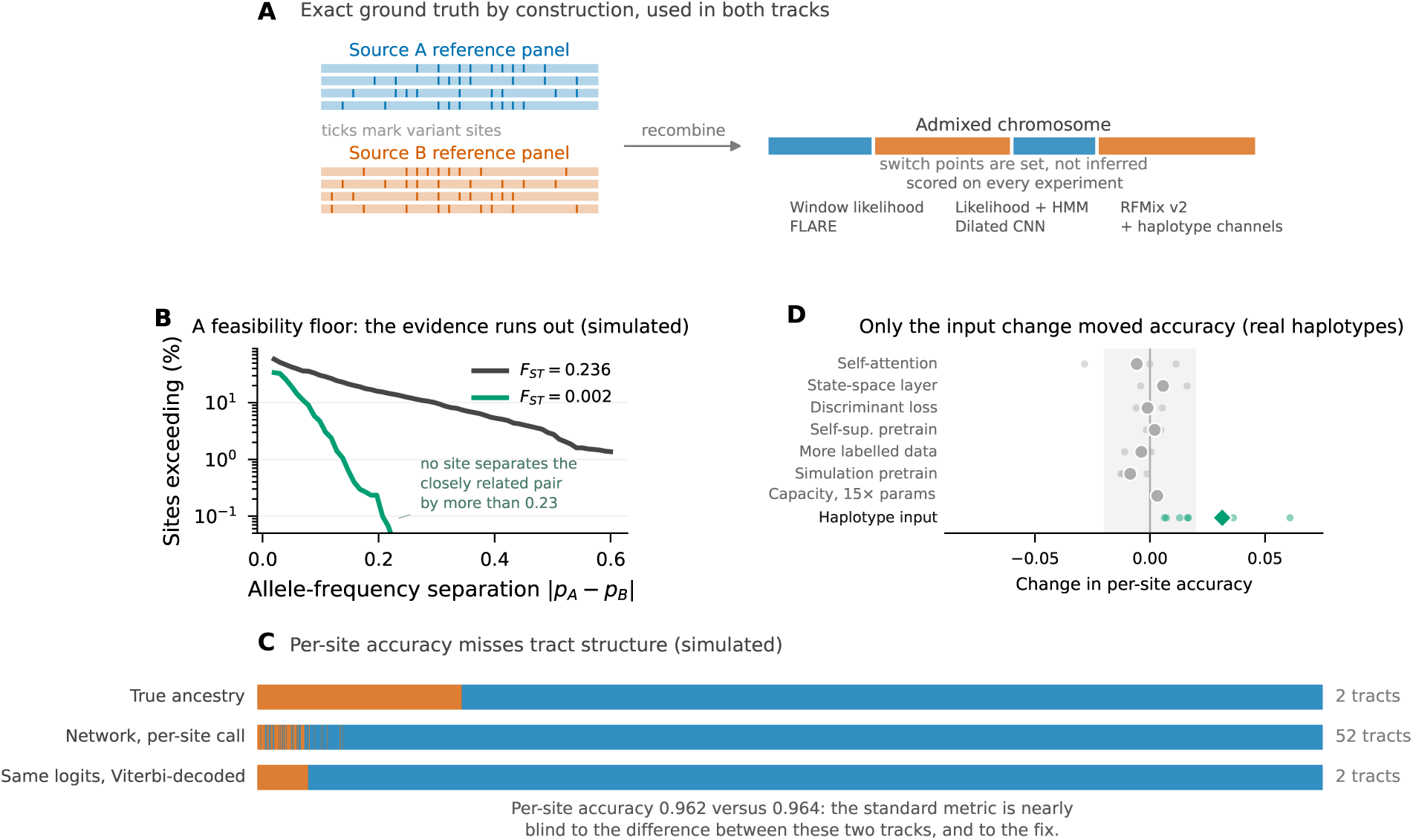
The task, the constraint, and what moved accuracy. (*A*) The construction, drawn schematically: haplotypes from two source reference panels are recombined into an admixed chromosome whose ancestry switch points are set rather than inferred, so every site carries an exact label. The same construction serves both the coalescent simulations and the real 1000 Genomes panels, and the six methods named are scored on identical sites throughout. (*B*) Percentage of sites at which the two source panels’ allele frequencies differ by more than the value on the horizontal axis, for simulated panels at the two ends of the divergence sweep: at *F_ST_* = 0.0020 no site separates the sources by more than 0.23, whereas at *F_ST_* = 0.236 15.5% of sites exceed 0.2. (*C*) A 2600-site window of one simulated admixed haplotype at *F_ST_*= 0.036: true ancestry, the network’s per-site calls, and the same logits after Viterbi decoding, at 0.962 and 0.964 per-site accuracy respectively. Per-site accuracy differs by +0.0002 between the two predicted tracks, which contain 52 and 2 ancestry tracts against a true 2. (*D*) Change in per-site accuracy on real haplotypes for every intervention we tried that was not a change of input, each against its own control and paired on seed, individual seeds behind the mean, shaded band *±*0.02; the input change is the diamond. Supplemental Table S3 gives the values, the controls, and the one arm omitted here. Panel A is a diagram; B, C and D are measurements, each title stating its data source.

## Results

### Accuracy declines steeply with source divergence

Fig. 2 shows per-site local-ancestry accuracy as a function of Hudson’s *F_ST_* between the two source populations, for all five methods. Each point is the mean over 512 held-out admixed haplotypes drawn from replicate simulations disjoint from those used for training; error bars give the standard deviation across replicates.

**Figure 2.**
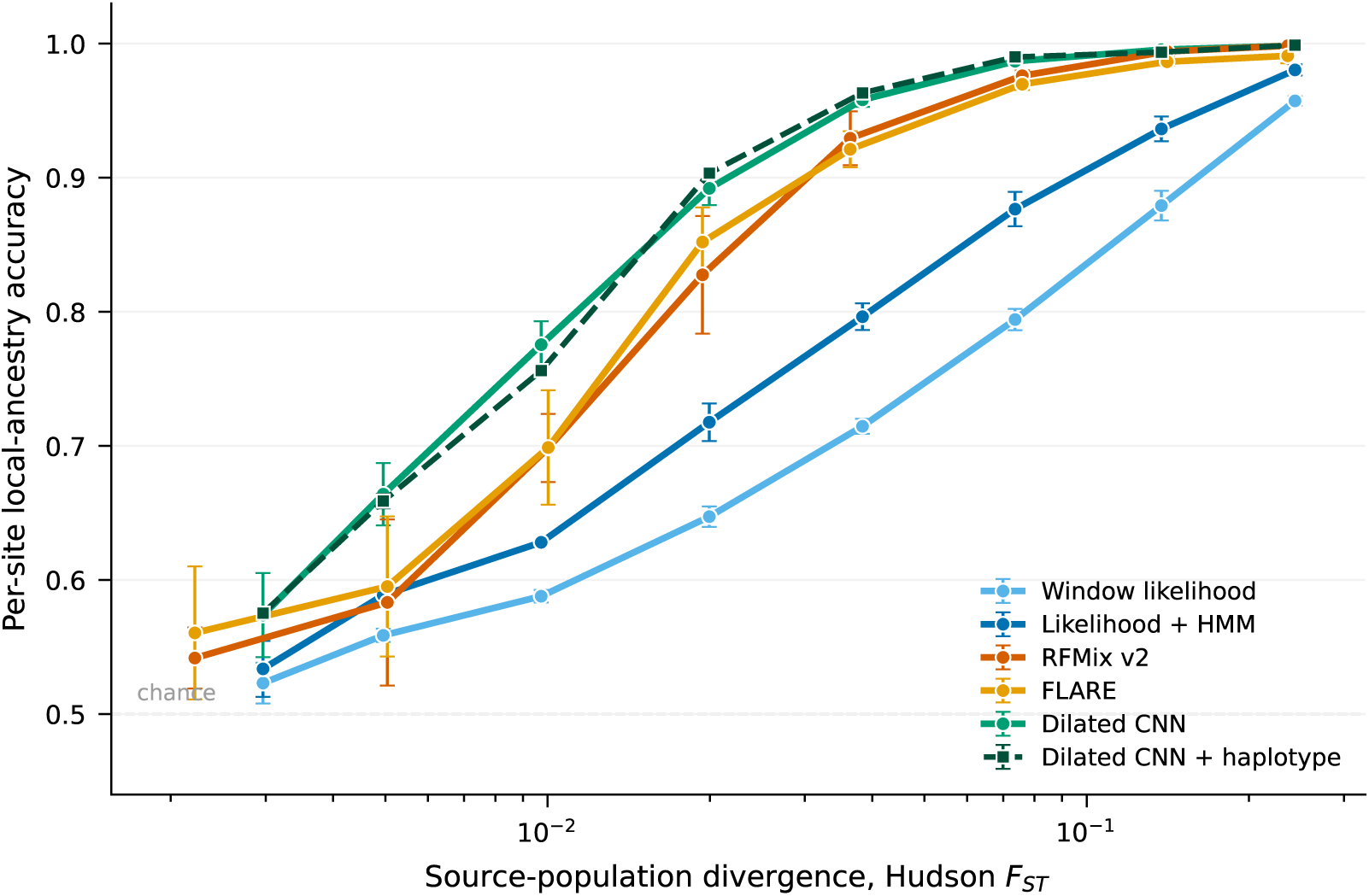
Local-ancestry accuracy as a function of source divergence. Per-site accuracy for five methods across 8 divergence levels. Points are means over held-out test replicates, error bars standard deviations across them. Dotted vertical lines mark two empirical *F_ST_* landmarks: the northern/southern Han comparison, and the European/East Asian comparison at which LAI methods are conventionally benchmarked. The dashed horizontal line is chance. The haplotype-aware network (dashed, squares) is the same architecture with four haplotype-matching channels appended, from an independent 5-seed replication plotted against the main sweep’s *F_ST_*by matched split time; because that replication varies only initialisation while the others vary the simulation, its bars span the narrower source of variation and are near-invisible here. The two curves nearly coincide: in simulation the channels are worth +0.003, in contrast to their effect on real haplotypes.

All methods degrade monotonically as the sources become less differentiated. The HMM-smoothed likelihood baseline falls from 0.980 *±* 0.004 at *F_ST_* = 0.2434 to 0.534 *±* 0.021 at *F_ST_* = 0.0030, a decline of 0.447. The unsmoothed windowed likelihood classifier is uniformly worse, confirming that spatial smoothing contributes substantially at every divergence level.

At the divergence closest to the northern/southern Han comparison (*F_ST_* = 0.0050), the windowed likelihood classifier reaches 0.559, the HMM-smoothed version 0.589 and the network 0.664. Only the last of these is far enough above chance to support downstream use; the divergence at which usable accuracy begins is therefore above, not at, the level of within-Han or within-Europe structure.

### In simulation, learned segmentation leads at low divergence

Because all methods are scored on the same simulated replicates, the difference between them is a paired quantity and is estimated far more precisely than the accuracies themselves: at *F_ST_* = 0.0100 the individual accuracies vary across seeds by *±*0.03 while their difference varies by *±*0.006 (*t* = 13.8 over 3 seeds). Pooling the levels below *F_ST_* = 0.015, the network leads in 8 of 9 runs with a mean advantage of 0.021. We stress this because single unpaired runs at low divergence proved badly misleading during development, returning gaps of either sign; the paired comparison is what makes the simulated advantage measurable at all. Against our own likelihood-plus-HMM baseline the same advantage measures 0.174, roughly four times larger and an artefact of that baseline’s weakness rather than a property of the network.

### Comparison with released implementations

A reimplemented baseline can flatter a new method, so we additionally ran RFMix v2 and FLARE on the identical simulated haplotypes, exported to VCF with matching genetic maps and with each tool given the true number of generations since admixture. The comparison is unforgiving of our own baseline: at *F_ST_* = 0.0364, the windowed-likelihood-plus-HMM baseline reaches 0.796 while RFMix reaches 0.929, a gap far larger than any difference between methods we would otherwise have reported. Conclusions drawn against our baseline alone would have substantially overstated the value of learned segmentation.

Against the released tools the learned network retains an advantage, but a much smaller and strongly divergence-dependent one. It peaks at 0.048 *±* 0.006 at *F_ST_* = 0.0100 (95% CI [+0.033, +0.063]; 0.765 versus 0.699 for the better of RFMix and FLARE), and falls to -0.004 at the lowest divergence tested and -0.000 at the highest, where all methods saturate and RFMix is marginally ahead. The peak accuracies do not subtract to the quoted gap, because the gap is computed per replicate and then averaged, and which of RFMix and FLARE is better varies between replicates; the paired figure is the conservative one and is what we report throughout. Fig. 2 shows all five methods on common axes.

### The trained network’s reliance on its long-dilation blocks is concentrated

If the learned method’s advantage comes from integrating weak per-site evidence over longer stretches of sequence than a fixed window spans, then the network’s reliance on its long-dilation blocks should track the advantage itself. We tested this by removing contiguous groups of dilated blocks from the trained networks. Because each block is residual, deleting it is exactly the identity map, so the network remains well-formed without retraining.

Single-block ablation is uninformative here and we do not report it: with residual connections the remaining blocks compensate for any one removal, so per-block importance is near zero throughout even where the blocks collectively matter. This is a general hazard of unit-level ablation in residual architectures and, we suggest, a reason to prefer group ablation when using pruning as a measurement rather than a compression tool.

Removing all blocks with dilation *≥* 8 leaves the fraction of above-chance accuracy reported in Fig. 3A. Retention falls to a minimum of 0.201 *±* 0.135 at *F_ST_* = 0.0200 and recovers to 0.815 *±* 0.076 at the highest divergence tested. At the two lowest divergences the ratio is not usefully estimable: its denominator, the above-chance accuracy of the unablated network, is below 0.20 there, and the spread across seeds (0.639 at *F_ST_* = 0.0030) exceeds the mean, so those points are shown open in Fig. 3A and we do not interpret them. The minimum falls inside the band of divergences where the network’s advantage over the better likelihood baseline is within 80% of its maximum. The trained network’s reliance on its long-dilation blocks is thus greatest in the same band in which the learned method’s advantage is greatest, and least where per-site evidence is abundant enough that the surviving blocks suffice. The linear discriminant criterion computed on the same networks (Fig. 3B) rises monotonically with divergence and collapses at the floor. Because *J* is measured on learned features, a low value is on its own consistent either with information being absent or with the network failing to extract it; read together with the failure of the likelihood baselines at the same divergences (Fig. 2), the first reading is the one the data support.

**Figure 3.**
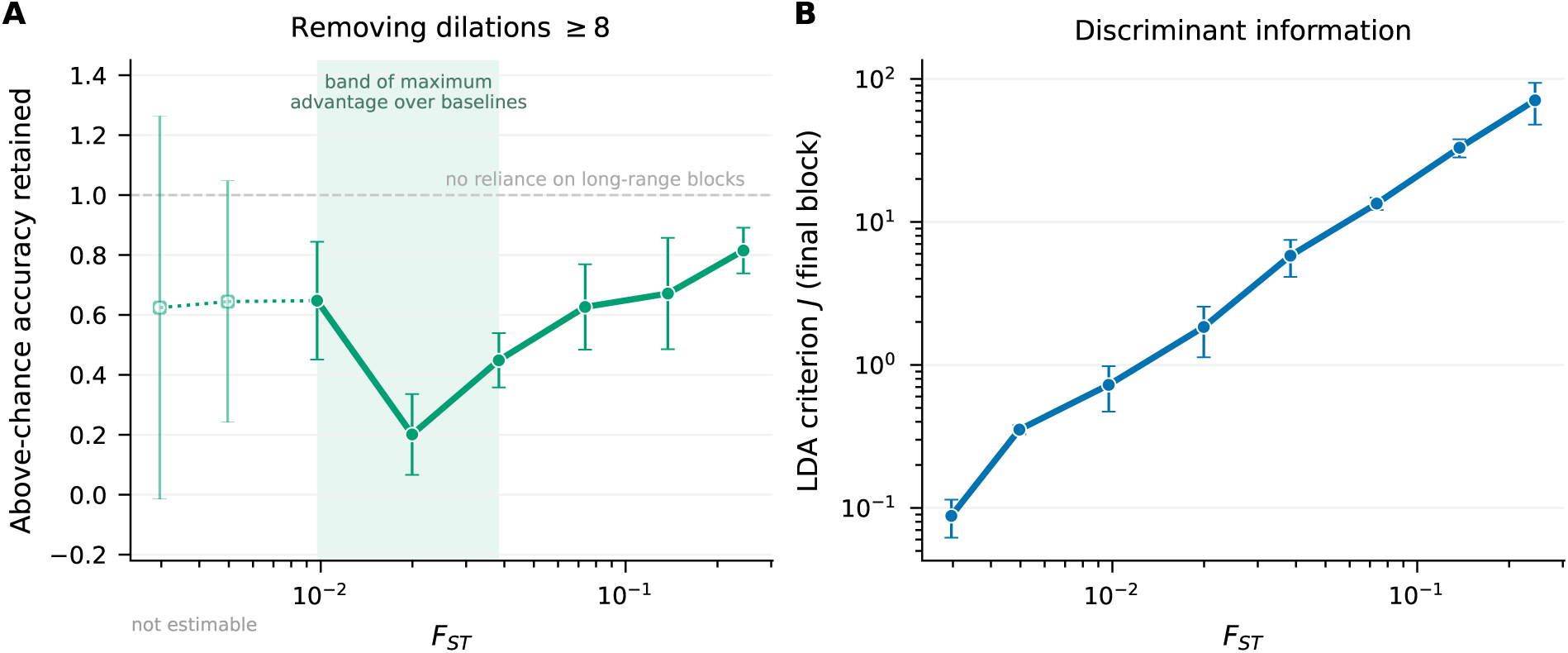
Reliance on long-range integration, and discriminant information. (*A*) Fraction of above-chance accuracy retained after removing all residual blocks with dilation *≥* 8. Low retention means the network depends on evidence integrated over long stretches of sequence; retention near unity (dashed line) means the surviving blocks suffice. Open symbols on a dotted segment mark divergences where the ratio is not usefully estimable, its denominator falling below 0.20; they are not interpreted. The shaded band spans the divergences at which the network’s advantage over the better likelihood baseline is within 80% of its maximum (Fig. 2); the retention minimum falls inside it. (*B*) Linear discriminant criterion *J* on the final-block features of the same networks, logarithmic axes. Points are means over 3 replicates with standard deviations.

### Sensitivity to divergence misspecification

Simulation-trained methods are commonly criticised on the grounds that they learn the simulator rather than the underlying biology. Supplemental Fig. S1 addresses this directly: every trained network is evaluated against every held-out test set, giving accuracy as a function of both training and evaluation divergence.

The matched-divergence diagonal averages 0.851. The largest off-diagonal degradation relative to the matched model is 0.169 (trained at *F_ST_* = 0.1374, evaluated at *F_ST_*= 0.0205). Misspecification is nevertheless mild in the region that matters: a single network trained at *F_ST_*= 0.0205 attains a mean of 0.850 across all evaluation divergences, within 0.001 of the 0.851 achieved by an oracle that always selects the matched specialist. Accurate prior knowledge of source divergence is therefore not required, and there is no ensemble of specialists worth combining.

### A feasibility floor below which no method is informative

The low-divergence end of Fig. 2 is qualitatively different from the rest of the curve. At *F_ST_* = 0.0022 the best of the five methods reaches 0.575, and the spread between the worst and best method is smaller than the seed-to-seed variability of any one of them. This is not a regime in which these methods differ in quality; it is one in which none of them extracts usable information from the data.

This matters because real population pairs fall below it. Measured on chromosome 22 of the 1000 Genomes phased release, CHB versus CHS gives *F_ST_* = 0.00042, IBS versus TSI 0.00152, and GBR versus CEU 0.00003, all beneath the floor. The implication is not that LAI is merely difficult for such pairs but that, with the methods and panels available here, it is uninformative, and that published inferences of fine-scale ancestry tracts within Han Chinese or within European populations require independent support.

### Per-site accuracy conceals a failure of tract structure

Per-site accuracy is the standard metric in this literature, but it scores each position independently and is therefore blind to whether the recovered mosaic has the right shape. This matters because several downstream uses read tract structure rather than per-site labels: admixture dating in particular infers the time since admixture from the tract-length distribution, so a method that systematically fragments tracts biases the inferred date however well it labels individual sites.

Scoring the same predictions by tract structure separates the methods far more sharply than accuracy does, and in the opposite direction (Table 1). Across 8 divergence levels RFMix recovers essentially the correct number of tracts (1.3*×* the true count, mean length 1.45*×* truth) and FLARE nearly so (1.0*×*, 1.62*×*). The convolutional network instead produces 78.8*×* too many tracts, rising to 209*×* at the lowest divergence, with mean tract length 0.30*×* truth. Our own likelihood baselines fragment similarly (22.6*×* and 6.6*×*).

**Table 1.** Tract-level structure, averaged over 8 divergence levels. Ratios are predicted to true. A tract-count ratio near one and a length ratio near one indicate a correctly shaped mosaic. Methods that fragment the mosaic can still score well on per-site accuracy. Tract-count ratios scale with the sequence they are measured over, so all rows are scored by identical tiling of the full replicate. Every row covers all 8 divergence levels.

| Method | Tracts (pred/true) | Mean length (pred/true) |
| --- | --- | --- |
| Window likelihood | 22.6 | 0.07 |
| Likelihood + HMM | 6.6 | 0.23 |
| RFMix v2 | 1.3 | 1.45 |
| FLARE | 1.0 | 1.62 |
| Dilated CNN (thresholded) | 78.8 | 0.30 |
| Dilated CNN (Viterbi-decoded) | 1.56 | 0.91 |
| Dilated CNN + haplotype (thresholded) | 42.6 | 0.36 |
| Dilated CNN + haplotype (Viterbi-decoded) | 1.90 | 0.81 |

The network therefore wins on per-site accuracy while destroying the structure the inference exists to recover, and no inspection of the accuracy curves would reveal it.

The consequence downstream can be quantified rather than asserted. Under a single-pulse model, ancestry tract lengths are approximately exponential with mean 1*/g* Morgans, so a method’s tracts imply an admixture time *g*^ = 1*/L̄*. Applied to the true tracts this estimator returns 25 generations against a simulated 30, a modest downward bias inherent to the estimator and to finite chromosome length; method bias is therefore best read relative to that value. RFMix implies 22 generations (0.9*×* the truth-derived value) and FLARE 25 (1.0*×*), both essentially correct. The convolutional network implies 1530 generations, a 61.2*×* overestimate, and our own likelihood baselines 128 and 433 (5.1*×* and 17.3*×*). A study dating this admixture from the network’s output would place a 30-generation event in the deep past.

The fragmentation is nevertheless a property of the decision rule rather than of the representation, and it is removable at no cost. The network emits an independent decision per site, with nothing penalising a switch; RFMix and FLARE cannot produce a shattered mosaic because each carries an explicit switching process. Imposing the same constraint on our network without retraining, by decoding the unchanged logits under a transition prior taken from the recombination process rather than learned, brings the tract-count ratio to 1.56*×*, the mean tract length to 0.91*×* truth, and the implied admixture time to 34 generations against 26 from the true tracts. This is a paired comparison of the two decodings of identical logits, across 8 divergence levels and 3 seeds (Table 1); the single-seed thresholded figures quoted above differ from its thresholded arm only by seed variation.

Per-site accuracy moves by +0.0002, that is, very slightly in the network’s favour, on 19 of 24 runs. This is the sharpest statement of the problem with the metric: it is blind to a defect that renders the output unusable for admixture dating, and equally blind to the fact that a change of decision rule costing nothing at all removes it. A study reporting only per-site accuracy would have shipped the fragmented output and never discovered that the improvement was available.

We caution against reading breakpoint-localisation error in isolation. The median distance from each true breakpoint to the nearest predicted one is smaller for the fragmenting methods, but only because predicting many spurious breakpoints places one near any given true breakpoint by chance. That statistic is interpretable only jointly with the tract-count ratio.

### On real haplotypes the learned method loses its lead

Coalescent simulation permits divergence to be varied continuously but imposes an idealised demography. We therefore repeated the comparison using phased 1000 Genomes haplotypes as source panels, holding the mosaicking construction and every downstream step fixed, across 11 population pairs spanning *F_ST_* from 0.00042 to 0.10912 and three superpopulations (Table 2, Fig. 4).

**Figure 4.**
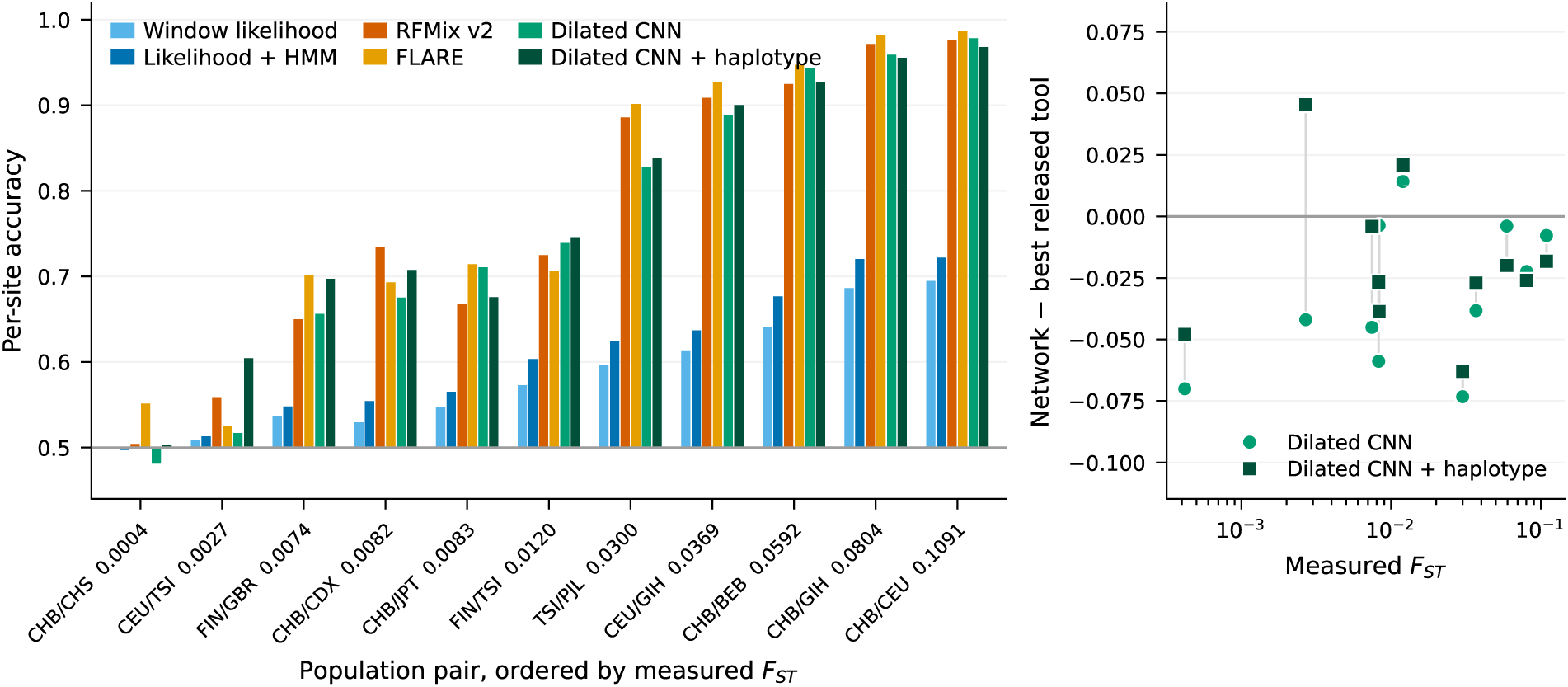
Real 1000 Genomes population pairs. Left: per-site accuracy for all six methods on 11 real pairs, one group of bars per pair, ordered by measured *F_ST_* and labelled with it. Bars are drawn from chance rather than from zero, since chance is the meaningful floor for a two-way call and a below-chance result then reads as a bar beneath the line; the horizontal line marks chance. Right: the difference between each network configuration and the better of the two released tools, against measured *F_ST_* on the same logarithmic axis as Fig. 2, with a pair’s two configurations joined by a thin line so the shift is readable per pair. Points are not joined across pairs, since these are discrete population pairs rather than a swept parameter, and pairs are labelled in Fig. 5 rather than here, where two points per pair would collide. With frequency input alone the network leads on 1 of 11 pairs and its deficit shows no monotone dependence on divergence (Spearman *ρ* = +0.45, *p* = 0.16).

**Table 2.** Per-site accuracy on real 1000 Genomes population pairs. Source panels are phased haplotypes from the named populations; admixed haplotypes are mosaicked from held-out donors so that ground truth remains exact. All methods receive 80 reference haplotypes here; Table 3 repeats the comparison at 92, and accuracies at both should be read as depending on that choice. Pairs span two orders of magnitude in divergence, six within and five across superpopulations; 1000 Genomes population codes are expanded in Methods. With frequency input alone the learned network is beaten by a released tool on 10 of 11 pairs. The final column adds haplotype-matching channels and is the mean of five seeds from the input-parity experiment; the preceding column is the single run of the original sweep, in the same harness.

| Pair | $F_{ST}$ | Window likelihood | Likelihood + HMM | RFMix v2 | FLARE | Dilated CNN | Dilated CNN + haplotype |
| --- | --- | --- | --- | --- | --- | --- | --- |
| CHB/CHS | 0.00042 | 0.498 | 0.497 | 0.504 | 0.551 | 0.481 | 0.503 |
| CEU/TSI | 0.00269 | 0.509 | 0.513 | 0.559 | 0.525 | 0.517 | 0.604 |
| FIN/GBR | 0.00745 | 0.536 | 0.548 | 0.650 | 0.701 | 0.656 | 0.697 |
| CHB/CDX | 0.00825 | 0.529 | 0.554 | 0.734 | 0.693 | 0.675 | 0.707 |
| CHB/JPT | 0.00831 | 0.547 | 0.565 | 0.667 | 0.714 | 0.711 | 0.676 |
| FIN/TSI | 0.01198 | 0.573 | 0.604 | 0.725 | 0.707 | 0.739 | 0.746 |
| TSI/PJL | 0.03002 | 0.597 | 0.625 | 0.886 | 0.902 | 0.828 | 0.839 |
| CEU/GIH | 0.03689 | 0.613 | 0.637 | 0.909 | 0.927 | 0.889 | 0.900 |
| CHB/BEB | 0.05922 | 0.641 | 0.677 | 0.925 | 0.947 | 0.943 | 0.927 |
| CHB/GIH | 0.08043 | 0.686 | 0.720 | 0.972 | 0.981 | 0.959 | 0.955 |
| CHB/CEU | 0.10912 | 0.695 | 0.722 | 0.977 | 0.986 | 0.978 | 0.968 |

**Table 3.**
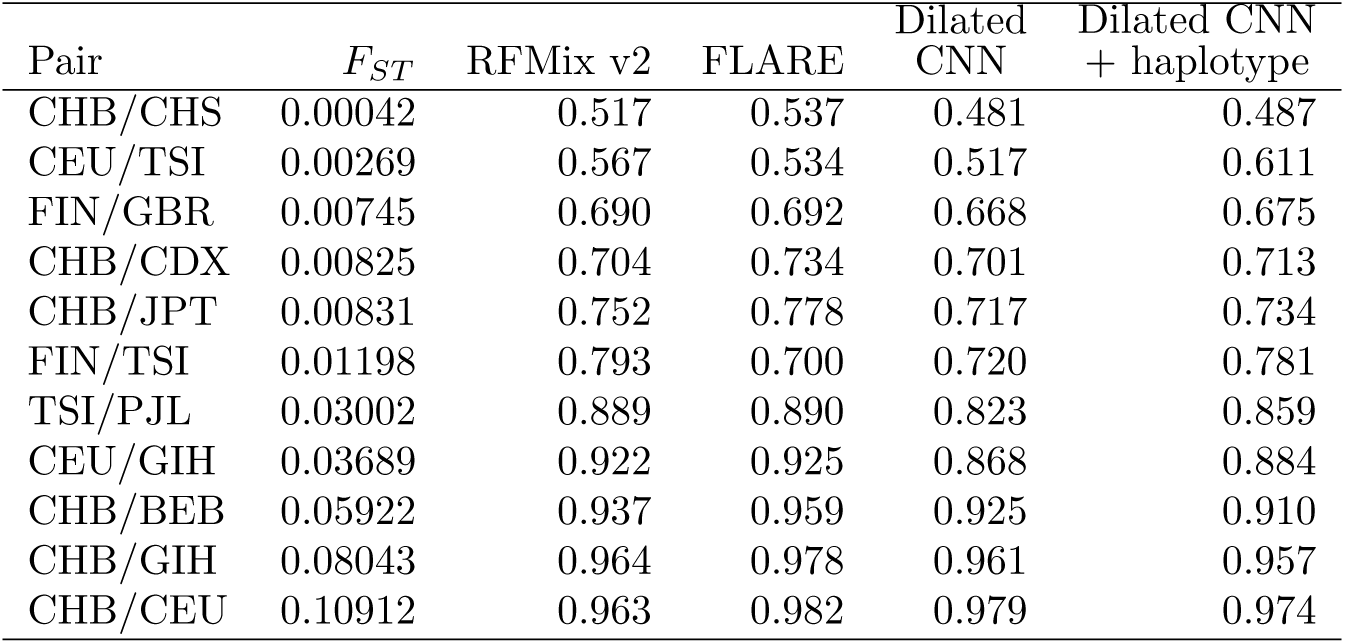
Per-site accuracy at the enlarged reference panel. All methods receive 92 reference haplotypes, the largest number common to every pair, with the donor set held at 80 so the evaluation data are identical across conditions. Network values are means of five seeds; the released tools are single runs, as in Table 2. Compare with Table 2, which uses 80 reference haplotypes: every method improves and their ordering is essentially unchanged. The haplotype-aware network exceeds the better released tool on 1 of 11 pairs.

| Pair | $F_{ST}$ | RFMix v2 | FLARE | Dilated CNN | Dilated CNN + haplotype |
| --- | --- | --- | --- | --- | --- |
| CHB/CHS | 0.00042 | 0.517 | 0.537 | 0.481 | 0.487 |
| CEU/TSI | 0.00269 | 0.567 | 0.534 | 0.517 | 0.611 |
| FIN/GBR | 0.00745 | 0.690 | 0.692 | 0.668 | 0.675 |
| CHB/CDX | 0.00825 | 0.704 | 0.734 | 0.701 | 0.713 |
| CHB/JPT | 0.00831 | 0.752 | 0.778 | 0.717 | 0.734 |
| FIN/TSI | 0.01198 | 0.793 | 0.700 | 0.720 | 0.781 |
| TSI/PJL | 0.03002 | 0.889 | 0.890 | 0.823 | 0.859 |
| CEU/GIH | 0.03689 | 0.922 | 0.925 | 0.868 | 0.884 |
| CHB/BEB | 0.05922 | 0.937 | 0.959 | 0.925 | 0.910 |
| CHB/GIH | 0.08043 | 0.964 | 0.978 | 0.961 | 0.957 |
| CHB/CEU | 0.10912 | 0.963 | 0.982 | 0.979 | 0.974 |

The floor reproduces. At *F_ST_* = 0.00042 (CHB/CHS) the best of the five methods reaches 0.551, and at *F_ST_* = 0.00269 (CEU/TSI) 0.559; both are within the range that separates methods from chance elsewhere in the benchmark. Local ancestry between these pairs was not recovered by any method we ran.

The simulated advantage does not reproduce. Where simulation placed the convolutional network ahead of the better released tool by up to 0.048, on real haplotypes it is beaten by a released tool on 10 of 11 pairs, with a mean deficit of -0.032 and a largest deficit of 0.073 (TSI/PJL). It leads on 1 pair (FIN/TSI). The deficit is not confined to the low-divergence regime: it appears at every level tested from *F_ST_*= 0.00042 to 0.08043. We place little weight on the highest-divergence pair, where all three of the stronger methods exceed 0.97 and the comparison is saturated, but the mid-range pairs carry the point on their own.

Our likelihood-plus-HMM baseline remains the weakest method on real data as it was in simulation, RFMix exceeding it on every pair, so the change is specific to the learned method: it is not that everything we implemented degrades, but that the one whose apparent advantage was established in simulation fails to retain it.

Divergence explains little of the variation between pairs. The deficit ranges from -0.073 to +0.014 with no statistically significant dependence on *F_ST_* (Spearman *ρ* = +0.45, *p* = 0.16; a weak positive tendency that eleven pairs cannot resolve), and two pairs at indistinguishable divergence (CHB/CDX and CHB/JPT, differing in *F_ST_* by under 1%) differ in deficit by 0.055. Source distinguishability is therefore not captured by *F_ST_* alone at this resolution, and the divergence axis should be read as an ordering rather than a sufficient statistic for difficulty.

The explanation lies in what the two families of method are given rather than in what the simulator generates. RFMix and FLARE both consume reference haplotypes; the network as specified above consumes allele frequencies and their likelihood ratio, which marginalise away which reference haplotype a segment resembles. A shortfall of haplotype content in the simulated panels does not account for the difference. Scoring ancestry by haplotype matching alone (the mean of the top-*k* agreement fractions against each panel, with no frequencies, no smoothing and no learning) gives higher accuracy on simulated panels than on real ones at matched divergence, on 4 of 5 pairs, by 0.048 on average (interpolating the simulated curve to each real pair’s *F_ST_* ; with five comparisons this does not reach significance). What the deficit reflects is the network’s input representation, and the following section closes it directly.

### Gene flow, not the recombination map, accounts for the simulated advantage

If the clean split is what flatters the learned method, then relaxing it should cost the method its advantage, and relaxing the other obvious simplification should not. We crossed continuous symmetric migration with an empirical recombination map in a 36-run factorial, three split times per condition and 3 seeds per cell (Supplemental Table S4). Migration reverses the advantage: the network’s margin over the better released tool is +0.033 under a clean split and -0.013 under migration. The empirical map does nothing, +0.034 against +0.033.

The raw condition means confound the comparison, because migration suppresses *F_ST_* : the migrating arms span 0.0026 to 0.0067 where the clean-split arms reach 0.0181, and the network’s advantage is itself divergence-dependent. Two controls separate them. Restricted to *F_ST_ <* 0.007, where all four conditions have runs, migrating arms give -0.009 against +0.028 without migration (Welch *p* = 0.002, *n* = 18 and 5). Regressing the gap on *F_ST_*, migration and map across all 36 runs gives a migration coefficient of -0.042 (SE = 0.011, *p <* 0.001) against +0.004 for the map (*p* = 0.672).

We report this as a bounded probe rather than a measurement. 3 seeds per cell establish the sign and the rough magnitude of the migration effect, not a precise estimate, and the divergence ranges the conditions span are not matched by construction. Read that way it identifies which simplification matters: gene flow, which real closely related populations have and a clean split does not, and not the recombination map that simulation studies more often worry about. It also answers the obvious question about our own design. We report a clean-split sweep because it is what the field simulates; that it inflates a learned method’s apparent advantage is part of what this paper is for.

### The deficit is one of input, and is partly recoverable below *F_ST_ ≈* 0.04

The preceding comparison is not input-matched: the released tools consume reference haplotypes, the network consumes panel frequencies. We therefore gave the network the same class of information and repeated the real-data experiment. Matching a query against reference haplotypes and learning on the result is not itself new: SALAI-Net computes a cosine similarity between query and reference windows, equivalent to a rescaled Hamming distance, and pools the top-*k* scores per ancestry (Oriol Sabat et al. 2022). The channels below are of that family. What the present experiment adds is the divergence range over which such information matters, and the demonstration that its absence, rather than any of the architectural choices we varied, is what the earlier comparison was measuring. That second point rests on testing the architectural explanations directly, which we did before changing the input, and which is bounded by the six interventions we could run: self-attention, a state-space layer, a fifteen-fold range of parameter counts, a discriminant training objective, self-supervised pretraining and more labelled data each move accuracy by at most 0.006, with inconsistent sign (Supplemental Table S3). Four haplotype-matching channels were appended to the four published ones (for each panel, the best and the mean top-*k* agreement fraction in a local window), leaving the architecture, optimiser, windows and partitions untouched. Parameter count rises by under 1%, so any change is attributable to the input rather than to capacity. Each configuration was trained under five seeds sharing identical partitions and windows, and is compared pairwise on seed.

The effect depends on divergence and changes sign (Fig. 5, left; Supplemental Table S1). Below *F_ST_* = 0.04 the haplotype-aware network gains +0.023 on average across 8 pairs, improving on 7 of them (one-sample *t* = 2.63, *p* = 0.034), with the largest gain +0.065 at CEU/TSI (*F_ST_* = 0.0027). Above that divergence it loses 0.014 on average and improves on 0 of 3 pairs. The two regimes differ (pooled two-sample *t* = 2.45, *p* = 0.037; sign pattern Fisher exact *p* = 0.024), and the gain declines with divergence across all 11 pairs (Spearman *ρ* = *−*0.58, *p* = 0.060). The interpretation is mechanical rather than mysterious: where allele frequencies barely separate the sources, haplotype identity carries information that frequencies do not; where divergence is ample, the frequency channels already suffice and four additional channels add variance without adding signal. The same channels are worth +0.003 in simulation across 8 divergence levels (below), so what they supply is not a property of the features in isolation but of what allele frequencies fail to supply on real chromosomes.

**Figure 5.**
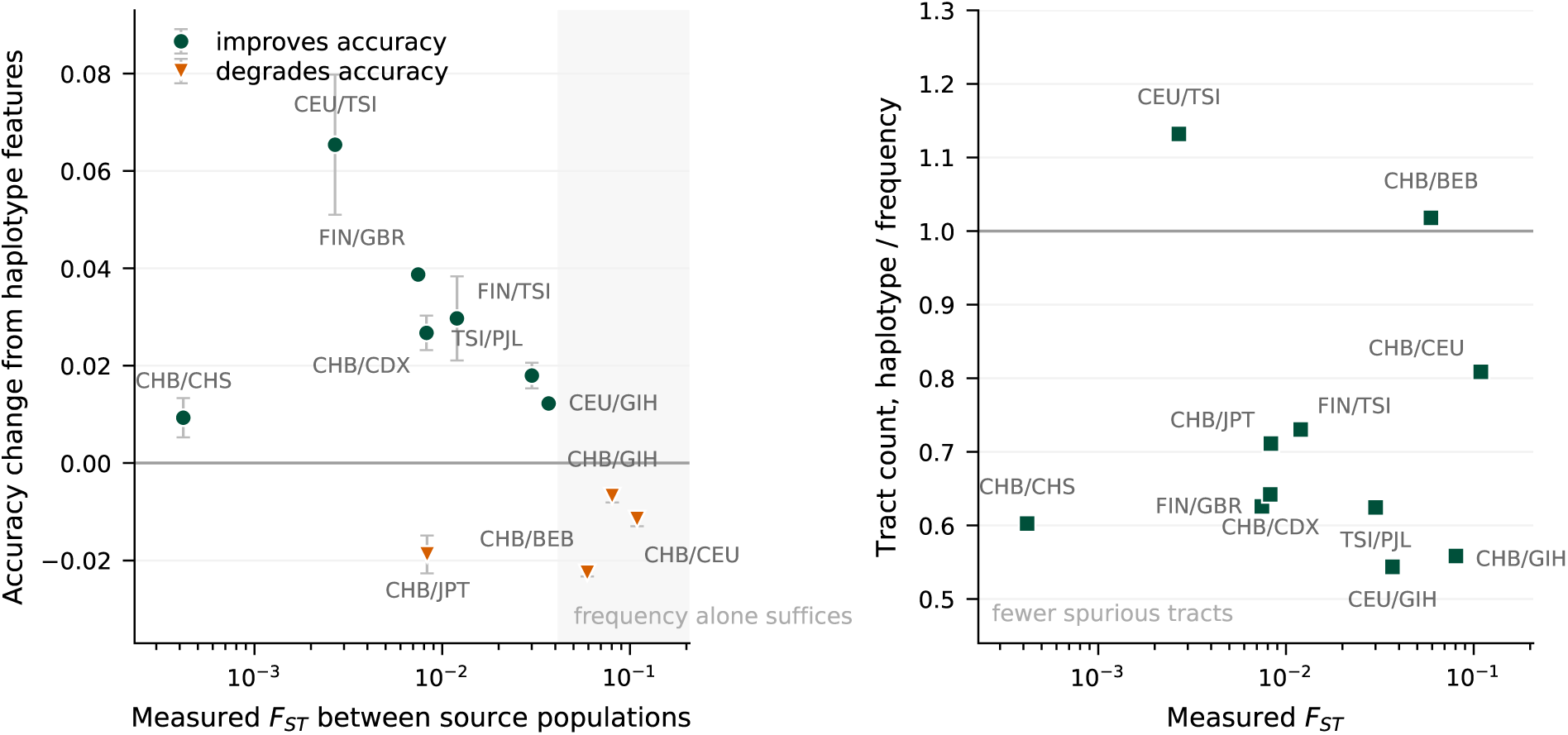
What haplotype features buy, against source divergence. Left: change in per-site accuracy from adding haplotype-matching channels, paired on seed, with bars giving the standard error over five seeds. Circles mark pairs that improve, triangles those that degrade; the shaded region is the regime above the estimated crossover, where allele frequencies alone already separate the sources. Right: tract count under haplotype input relative to frequency input, so values below one mean fewer spurious ancestry tracts. The two panels disagree by design: accuracy changes sign with divergence while tract structure improves on 9 of 11 pairs, which is why both are reported. Note that CEU/TSI, the largest accuracy gain, is the one low-divergence pair whose tract count worsens.

The change does not, however, make the network competitive with the released tools. Repeating the comparison with the reference panel enlarged to 92 haplotypes, the largest common to all 11 pairs, and with donors held fixed so that the evaluation data do not move, the effect on the network is larger and more consistent than at the smaller panel (+0.031 on 8 of 8 pairs below *F_ST_* = 0.04, *p* = 0.026; largest at CEU/TSI, +0.095). Yet the network exceeds the better released tool on 1 of 11 pairs, against 0 for the frequency-only configuration. Supplying the missing information improves the learned method substantially and still leaves it behind.

Where it does win is where the account predicts. The one pair on which the haplotype-aware network exceeds both released tools is CEU/TSI, at *F_ST_* = 0.0027, the least divergent pair at which any method is informative, the 1 pair below it reaching only 0.537 for any method. There it attains 0.611 against 0.567 for RFMix, a margin of 0.044, and the seed spread is 0.002, so the ordering is not a matter of initialisation. The same pair carries the largest gain from the channels themselves (0.517 for the frequency-only configuration, a gain of +0.095). The advantage the network holds is therefore narrow but not accidental: it is concentrated at the bottom of the usable range, exactly where allele frequencies are least able to separate the sources and haplotype identity carries what they cannot, and it is lost as soon as divergence is sufficient for frequencies to suffice. Tract structure improves more broadly than accuracy does (Fig. 5, right), on 9 of 11 pairs overall and by -22.9 tracts on average below the crossover, relevant because admixture dating reads the tract-length distribution rather than the per-site call.

Neither of the two remaining possibilities improves on this. Adding the positional channel on top of the haplotype channels changes accuracy by -0.006 across 5 pairs and helps on 0 of them; four of those pairs are ones on which the haplotype-aware network trails FLARE, and on none of them does it close the gap or overtake a released tool. The same channel alone was also null at CHB/CDX. Distance is not the missing information. And the two interventions that do help overlap rather than compound: haplotype channels reduce raw fragmentation from 78.8 to 42.6 tracts per true tract, but after Viterbi decoding the frequency-only network is the better shaped of the two (1.56 against 1.90), while the haplotype-aware network retains the higher accuracy (0.851 against 0.840). Decoding and input each supply what the other largely already provides.

The gain is visible in the representation and not only in the decision rule. Measuring the LDA criterion *J* = tr((*S_W_* + *λI*)*^−^*^1^*S_B_*) on the final block, haplotype input raises it on 8 of 8 divergence levels, by a factor of 1.77 on average. Absolute *J* varies by threefold across evaluation-window draws and is not reportable alone; the ratio between configurations on identical windows is stable to 0.31 and is what we report. Effective discriminant dimensionality does not move consistently, so the channels supply more separable information rather than recruiting more of the representation.

The same features do nothing in simulation, which is the sharper half of the comparison. In a matched ablation holding simulations, windows and optimiser fixed and repeating each configuration under 5 seeds, they change accuracy by +0.003 across 8 divergence levels, improving on 5 of them (*t* = 0.91, *p* = 0.39). Within a level the seed spread is small enough that individual differences are themselves significant; it is their sign that is inconsistent along the divergence axis. The contrast is not explained by simulated panels carrying less haplotype signal (measured above, they carry more) but by what the frequency channels substitute for in each setting. In simulation the frequency-only network already reaches 0.769 at *F_ST_ ≈* 0.01; averaged over the 4 real pairs between *F_ST_*= 0.005 and 0.015 it reaches only 0.687. There is a shortfall on real chromosomes for haplotype features to close, and none in simulation. A comparison conducted only in simulation would have concluded that this information is unnecessary.

Two limits should be read with these numbers. The crossover near *F_ST_* = 0.04 is estimated from these pairs rather than specified in advance, and only 3 pairs lie above it, so it locates a transition rather than validating a threshold. And one pair below the crossover, CHB/JPT, loses 0.019 at a divergence where a neighbouring pair gains +0.027, which is the same failure of *F_ST_* to determine difficulty reported above.

### Contiguity substitutes for a small reference panel

The haplotype channels above summarise, in a local window, the *rate* at which the target agrees with each reference haplotype. That is the statistic neural methods of this kind use (SALAI-Net computes a cosine similarity equivalent to a rescaled Hamming distance (Oriol Sabat et al. 2022)), and it is not the statistic the copying models use. RFMix and FLARE descend from Li–Stephens, whose sufficient quantity is the *length* of contiguous agreement, indexed efficiently by the positional Burrows–Wheeler transform (Durbin 2014). A reference agreeing at most sites at random and one matching perfectly across a long run are nearly identical in rate and wholly different as evidence of shared ancestry.

Measured as single-feature classifiers on the same data, with no learning and no smoothing, contiguity is the more discriminative of the two on 5 of 6 pairs, by +0.040 on average. Supplied to the network as additional channels it helps, but only where the reference panel is small: it adds +0.030 at 40 reference haplotypes, +0.023 at 80, and +0.003 at 100. The benefit decays as the panel grows, which is the opposite of what the mechanism naively suggests and is explained by the alternative rather than by contiguity itself: with few references the agreement rate is estimated from few samples and is noisy, while with many it becomes reliable and contiguity becomes redundant. Contiguity is therefore a substitute for reference panel size rather than an independent source of signal.

### Every method is far from reference-panel saturation

Reference panel size is treated as fixed in most comparisons, including ours until this point. It is not innocuous. Enlarging the panel from 80 to 100 haplotypes, with donors held fixed so the evaluation data are unchanged, raises accuracy for every method in every pair: +0.023 for the network, +0.028 for RFMix and +0.011 for FLARE, averaged over 4 pairs (Supplemental Table S2). The direction is the solid part of this result, all three methods improving in all 4 pairs, while the magnitudes are imprecisely estimated from so few pairs: the intervals are [+0.017, +0.030] for the network, [-0.003, +0.059] for RFMix and [-0.001, +0.023] for FLARE, and only the first excludes zero. For the network, where we also ran 40 reference haplotypes, the rise is monotone from 40 through 80 to 100 in every pair, with no sign of flattening. We therefore read this as evidence that none of the three is near saturation, and not as a measurement of how much each would gain, and the ordering between methods is essentially unchanged, so it is a property of the problem rather than an advantage available to any one method. It does mean that accuracies reported at a fixed panel size understate what each method can achieve, and that a comparison run at a single panel size cannot be assumed to hold at another.

### Phasing error costs integration, not input type

The comparison so far assumes perfectly phased haplotypes, which applied analyses never have. Switch errors have an informative property: exchanging alleles between the two haplotypes of one individual leaves the allele frequency spectrum exactly unchanged, so a method reading only frequencies cannot be affected, whereas reference haplotypes become chimeric and target haplotypes acquire ancestry transitions that no recombination produced. We therefore injected switch errors into reference panels and targets alike, at rates spanning good to poor phasing, training the network on data carrying the same error rate, the setting most favourable to it, since it can adapt to the noise rather than meeting it unseen (Table 4).

**Table 4.** Per-site accuracy under phase switch errors. Errors are applied to reference panels and targets alike; the network is trained at the same error rate it is tested at. The windowed likelihood classifier is unaffected by construction, since exchanging alleles within an individual leaves allele frequencies unchanged.

| $F_{ST}$ | Switches<br>per Mb | Window<br>likelihood | Likelihood<br>+ HMM | RFMix v2 | FLARE | Dilated<br>CNN | Dilated CNN<br>+ haplotype |
| --- | --- | --- | --- | --- | --- | --- | --- |
| 0.0105 | 0 | 0.593 | 0.634 | 0.693 | 0.748 | 0.770 | 0.762 |
| 0.0105 | 0.5 | 0.593 | 0.627 | 0.699 | 0.725 | 0.718 | 0.722 |
| 0.0105 | 2.0 | 0.588 | 0.617 | 0.665 | 0.713 | 0.680 | 0.707 |
| 0.0364 | 0 | 0.704 | 0.780 | 0.947 | 0.926 | 0.953 | 0.968 |
| 0.0364 | 0.5 | 0.701 | 0.771 | 0.883 | 0.899 | 0.923 | 0.948 |
| 0.0364 | 2.0 | 0.696 | 0.756 | 0.826 | 0.848 | 0.874 | 0.915 |

The invariance argument holds exactly. The windowed likelihood classifier changes by -0.005 at *F_ST_* = 0.0105 and -0.007 at *F_ST_* = 0.0364, that is, nothing.

Every other method degrades, and the frequency-only network degrades in a divergence-dependent way. At *F_ST_* = 0.0105 it is the most damaged of all methods, changing by -0.090 against -0.028 for RFMix and -0.035 for FLARE, and its advantage disappears entirely: at 0.5 switch errors per Mb, a rate consistent with good phasing, FLARE (0.725) overtakes it (0.718). At *F_ST_* = 0.0364 the ordering of sensitivities reverses, RFMix changing by -0.121 against -0.079 for the frequency-only network, which retains its lead at every rate tested.

Sensitivity is therefore not a property of reading frequencies or haplotypes, but of how far evidence is integrated. The windowed likelihood classifier reads frequencies and makes no use of context, and is unaffected. The network reads the same frequencies and integrates them over two thousand sites, and is the most damaged method in the study. Switch errors leave the frequency spectrum untouched while fragmenting exactly the tracts that integration exploits, so the method whose advantage rests on integration has the most to lose.

Supplying haplotype-matching channels reduces that exposure rather than increasing it, which is the opposite of what their reliance on contiguous segments would suggest. The haplotype-aware network changes by -0.055 and -0.053 across the two divergence levels, against -0.090 and -0.079 for the frequency-only network, and less than either released tool (-0.121 for RFMix and -0.077 for FLARE at *F_ST_* = 0.0364). At the highest error rate tested it is ahead of the frequency-only network by 0.026 and 0.041. The matching channels are pooled over local windows, which degrades gracefully as switches accumulate, whereas long-range integration does not. The configuration we recommend is thus the more robust of the two to the error that applied analyses actually carry, though we emphasise these are single runs per cell, and the effect does not extend to higher divergence.

## Discussion

Several conclusions follow from this benchmark, and they are of different kinds.

The first concerns the data. Below *F_ST_ ≈* 0.0022 none of the five methods carries usable information, in simulation or on real haplotypes, and the differences between them there are smaller than their own run-to-run variability. Within the settings we evaluated this behaves as a property of the genomes rather than of any one algorithm: the per-site and haplotype-level differences that local ancestry inference must exploit are very small. We state it as an empirical floor rather than a universal one, and two of our own results show it is not fixed. Supplying haplotype information lowers it: at *F_ST_* = 0.0027 the haplotype-aware network reaches 0.611 where the frequency-only configuration reaches 0.517. And it rests on a clean-split demography, which the factorial above shows is the condition most favourable to a learned method; what would move it is the amount of haplotype structure the sources carry, which continuous migration, unequal admixture proportions and multiple pulses all change. What we can say is bounded accordingly: no method we tested clears the floor under the conditions we tested. That is enough for the practical point, since the population pairs that most often motivate fine-scale ancestry analysis fall below it, northern versus southern Han Chinese by a wide margin, and claims of ancestry tracts within such populations therefore require support independent of the inference itself. Whether a future method could extract more from these data is not something our experiments can settle.

The second concerns what is measured. Per-site accuracy scores each position independently and is blind to the shape of the recovered mosaic. Judged by it our network is the best method in simulation; judged by tract structure it is among the worst, and both statements describe the same predictions. Since admixture dating and related analyses consume the tract-length distribution rather than per-site labels, the metric that dominates method comparisons in this literature is not the one that determines downstream usefulness. The point is sharpened by the fact that the fragmentation is removable by a change of decision rule that per-site accuracy cannot see. The metric is blind both to the defect and to its remedy, so a study guided by it would have shipped the broken output with the fix available and unnoticed.

The third concerns how methods are evaluated. Our convolutional network exceeded the better released tool by up to 0.048 in simulation, and was beaten by a released tool on 10 of 11 real population pairs, across two orders of magnitude of divergence. Had we validated on simulation alone, as is standard in this literature and as we initially intended, we would have reported an advantage that does not exist. The mechanism is not the one we first supposed. We expected a clean split to generate less haplotype sharing than real chromosomes carry, penalising methods that model reference haplotypes; measured directly, simulated panels carry more exploitable haplotype signal than real ones at matched divergence. What differs is what the frequency channels substitute for. In simulation the frequency-only network already reaches 0.769 at *F_ST_ ≈* 0.01, leaving nothing for haplotype information to add; on real pairs at the same divergence it reaches 0.687, and the shortfall is real. Our simulator flattered our own method, and no amount of replication within the simulation would have revealed it. Three independent seeds gave tight error bars around the wrong answer.

A fourth conclusion concerns what the shortfall consists of. We eliminated the architectural explanations we were able to test, one at a time, as reported above: none of the six interventions moved accuracy by more than 0.006, and their signs were inconsistent. One arm is a genuine exception and we report it as such: pretraining on simulation is worth +0.061 to the frequency-only network on 3/3 seeds, but *−*0.009 to the haplotype-aware one, which is what the domain-shift argument above predicts: the haplotype channels are computed against reference panels, and simulated panels are precisely where those panels differ most from real ones. The benefit is therefore an artefact of the weaker input rather than a route to closing the deficit, and it disappears once the input is fixed. Apart from that arm, the one intervention that mattered was changing what the network is shown. Supplying the haplotype-matching information the released tools already use recovers +0.031 (95% CI [+0.005, +0.058]) on 8 of 8 pairs below *F_ST_* = 0.04 and costs accuracy above it, where allele frequencies already separate the sources. The deficit is therefore representational more than it is a property of learned inference — but closing it is not sufficient, and the network still trails the better released tool on 10 of 11 pairs. The exception is informative rather than incidental: the pair it wins, CEU/TSI at *F_ST_*= 0.0027, is the least divergent one at which any method is informative, and it wins there by 0.044. A learned method given haplotype information is at its most useful precisely where the sources are hardest to tell apart, which is the regime this paper is about. What remains unexplained is the residual, and we do not attribute all of it to input. The copying models treat recombination explicitly where a convolution approximates it only through learned integration, and features of the simulation framework itself may contribute. Demographic misspecification we can now speak to: crossing continuous migration with an empirical recombination map shows that gene flow reverses the network’s simulated advantage while the map does not, so the demography that produced the training data is part of the account rather than a candidate we left untested. Our evidence separates input from architecture and from these two demographic factors; it does not separate it from the remainder.

Two quantities that comparisons conventionally hold fixed turned out to matter more than any architectural choice we tested. The first is the statistic used to summarise reference matching. Neural methods of this kind, ours included, use an agreement rate within a window; the copying models they are compared against use the length of contiguous agreement, which is the sufficient statistic of the underlying process. Contiguity is the more discriminative of the two as a raw feature, but supplying it to the network helps only where the reference panel is small (+0.030 at 40 reference haplotypes falling to +0.003 at 100) because the alternative improves as the panel grows rather than because contiguity degrades. It substitutes for panel size rather than adding to it. The second is panel size itself, which no method approaches saturating: enlarging it raises accuracy for the network and both released tools alike, without reordering them. Neither is an advantage available to any one method, and both mean that an accuracy reported at a single panel size understates what a method can do and cannot be assumed to transfer.

We draw a fifth methodological point from the comparison against our own baseline. Our likelihood-plus-HMM baseline, written in good faith to represent the classical approach, is far weaker than RFMix at moderate divergence, and against it the network’s apparent advantage was roughly four times larger than the one that survived contact with the released tools. Reimplemented baselines and simulation-only validation are two failure modes with the same signature: both inflate a new method’s apparent value, and both are invisible from inside the evaluation that produced them.

A sixth observation narrows the account further, though only partly. Applied LAI runs on statistically phased data, and phase switch errors leave allele frequencies exactly unchanged while fragmenting haplotypes. This costs the frequency-only network more than any other method, more than either released tool, which is initially surprising, since it reads only frequencies and switch errors leave frequencies unchanged. Sensitivity tracks how far evidence is integrated, not which input is read: the windowed likelihood classifier reads the same frequencies without context and is unaffected. Supplying haplotype channels reduces the exposure rather than increasing it, leaving the recommended configuration less sensitive than RFMix or FLARE. Phasing error is therefore a plausible contributor to the real-data result rather than a complete explanation of it, and we report it as such.

The mechanistic analyses support rather than drive these conclusions. Block-group ablation places the trained network’s greatest reliance on its long-dilation blocks in the same divergence band where its advantage is greatest. The linear discriminant criterion computed on the same representations speaks to a different point: it rises monotonically with divergence and is near zero at the lowest levels, corroborating that the floor reflects absent class-separating information rather than a failure to exploit information that is present. A further negative result is worth recording alongside them: cross-divergence transfer is mild enough that a single network trained at intermediate divergence performs within 0.001 of an oracle that always selects the matched specialist, so there is no ensemble worth distilling and no requirement to know source divergence in advance.

## Limitations

Several limitations bound the scope of these results, and we state them plainly because the benchmark is intended as practical guidance rather than as a methods advertisement.

First, the demographic and admixture models are idealised. Sources diverge cleanly from a common ancestor with no subsequent gene flow and constant effective population size, and we simulate a single pulse with equal contributions. Real closely-related pairs, northern and southern Han being a case in point, are separated by continuous gene flow rather than a clean split, and continuous or multi-pulse admixture with unequal proportions would change the tract-length distribution and shift absolute accuracies. Our real-data results bear the first point out directly: pairs at indistinguishable *F_ST_*differ in outcome by as much as 0.055, so the axis should be read as an ordering of source distinguishability rather than a sufficient statistic for difficulty. Which aspect of the demography matters we did test, and report above: gene flow reverses the network’s simulated advantage where an empirical recombination map does not. That probe is itself limited to 3 seeds per cell and to conditions whose divergence ranges are not matched, so it locates the responsible simplification without quantifying it.

Second, our phasing-error analysis uses a single run per cell and two divergence levels, and applies switch errors at a uniform rate rather than concentrated in regions of low information as real phasing error is. It also grants the network training data carrying the same error rate as the test data, which is more favourable than the applied setting, where a model trained on simulations meets real phasing error unseen. We also assume reference panel membership is known without error.

Third, the real-data arm remains semi-synthetic, in the sense set out in the Conclusion: the 1000 Genomes pairs supply real haplotypes, real linkage disequilibrium and a real allele frequency spectrum, but the admixture pulse is imposed rather than historical and the populations are sampling locations rather than the unadmixed sources of a documented admixture event. The loss of advantage we report is therefore a statement about haplotype structure rather than about admixture history.

Fourth, reference panel size is varied over a narrow range. The pairs admit at most 92 reference haplotypes if the donor set is to be held at 80 and identical across conditions, so we compare 40, 80 and 100 rather than the full panels an applied analysis would use. The direction is unambiguous, every method improving and none near saturation, but the shape of the curve beyond 100, and whether the methods separate at panel sizes we could not reach, are open. Our accuracies should accordingly be read as lower bounds that depend on a parameter most comparisons, including our own earlier sections, fix without comment.

Fifth, the divergence at which haplotype features stop helping is estimated from the pairs we ran rather than specified in advance. The sign of the effect changes between *F_ST_* = 0.037 and *F_ST_* = 0.059, and any cutoff in that interval yields the statistics we report, but only 3 pairs lie above it and all three are pairs on which every method already exceeds 0.92. The crossover should therefore be read as locating a transition in our data, not as a calibrated or transferable threshold. Whether it tracks divergence itself or the accuracy already attainable without the features is not resolved by 11 pairs, since the two are confounded across our panel.

Local ancestry inference has a divergence below which it recovers nothing, and several population pairs that motivate fine-scale ancestry analysis (northern versus southern Han Chinese, Iberian versus Tuscan, British versus Utah-European) lie beneath it. For those pairs, under every condition we evaluated, the limitation is the information content of the data rather than the sophistication of the method. We do not claim this as a universal bound: a demography that leaves more haplotype structure than a clean split would move the floor, and supplying haplotype information already moves it a little. We claim that it is where the floor sits under the conditions the field currently simulates.

One qualification governs the rest. Our real-data arm substitutes real haplotypes into a construction whose breakpoints we impose, because admixed individuals with known local ancestry do not exist at this divergence. It therefore tests how methods respond to real haplotype structure, real linkage disequilibrium and a real frequency spectrum, which is what the simulation-to-reality reversal turns on, and not how they perform on a naturally admixed cohort. Statements about what happens “on real haplotypes” should be read in that sense throughout.

Above the floor, three ways an evaluation can be internally consistent, well replicated and still misleading recur in what we found. The predictions that make our network the most accurate per site make it among the worst at recovering tract structure, which is what admixture dating consumes. Its simulated advantage does not survive contact with real haplotypes. And measured against a baseline we wrote ourselves, that advantage appeared several times larger than it does against the released tools.

What the learned method lacked was not depth but information. Haplotype-matching summaries recover most of the deficit where the sources are closely related, are superfluous once they are not, and leave the method ahead on the hardest comparison we ran and behind on the rest, a conditional and bounded statement, and the one the evidence supports. Two quantities that comparisons conventionally hold fixed, the statistic used to summarise reference matching and the size of the reference panel, moved accuracy more than any architectural choice we tested.

We therefore suggest five things of anyone comparing local ancestry methods: estimate *F_ST_* between the source panels before beginning, since it bounds what any method can achieve; report tract-level statistics alongside per-site accuracy, since the two can rank methods oppositely and since a decoding change invisible to one can transform the other; test under realistic phase switch error, which costs frequency-based and haplotype-based methods very differently; validate on real haplotypes against released implementations, since neither simulation nor a reimplemented baseline reliably predicts that comparison, and since *F_ST_* alone does not predict which real pairs prove hard; and, before attributing a learned method’s shortfall to its architecture, check whether it is receiving the same information as the tools it is being compared against. We would go further on the second of these: tract-level statistics should be reported as a matter of course rather than on request, since a method can be ranked first on per-site accuracy and last on the quantity a downstream analysis consumes. Breakpoint-localisation error and probability calibration are natural companions, though the first is interpretable only jointly with the tract count, as we show above, and we have not evaluated the second. None of these is expensive. All were necessary here to avoid reporting a result that was not true.

## Methods

### Coalescent simulation

Two source populations, *A* and *B*, descend from a common ancestral population of effective size *N_e_* = 10,000 diploids, splitting *T* generations before the present with no subsequent migration. We simulate 10 Mb of sequence under the Hudson coalescent with recombination using msprime 1.3.4 (Baumdicker et al. 2022), with per-base per-generation recombination rate *ρ* = 10*^−^*^8^ and mutation rate *µ* = 1.25 *×* 10*^−^*^8^, sampling haploid genomes. Divergence is varied by setting *T ∈ {*25, 50, 100, 200, 400, 800, 1600, 3200*}* generations, yielding 8 divergence levels spanning *F_ST_* from 0.0022 to 0.243.

Divergence is quantified by Hudson’s *F_ST_* estimator computed as a ratio of averages over segregating sites (Bhatia et al. 2013),

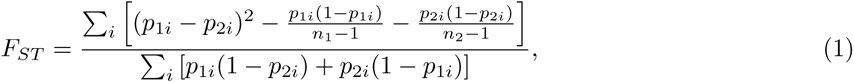

where *p_ji_* is the sample allele frequency in population *j* at site *i* and *n_j_* the number of sampled haplotypes.

From each population we draw 200 haplotypes, partitioned into a reference panel of 100 and a disjoint donor pool of 100. The disjointness matters: a donor haplotype never appears in the panel used to classify it, so the reported accuracies are not inflated by self-matching.

### Admixture construction and ground truth

Admixed haplotypes are constructed by mosaicking donor haplotypes rather than by simulating the admixture pulse coalescently. For each admixed haplotype we draw the number of crossovers from a Poisson distribution with mean *gρL*, where *g* = 30 generations since the pulse and *L* the sequence length, place breakpoints uniformly along the sequence, and assign each resulting segment independently to source *A* with probability 0.5, copying from a uniformly chosen donor haplotype of that source.

This construction yields local ancestry labels that are exact by construction rather than inferred from a tree sequence, so the ground truth against which all methods are scored is independent of any modelling choice.

### Methods compared

The three methods we implemented ourselves consume the target haplotype together with the allele frequencies of the two reference panels, clipped to [10*^−^*^3^, 1 *−* 10*^−^*^3^]. The per-site log-likelihood ratio for a target allele *x_i_*is

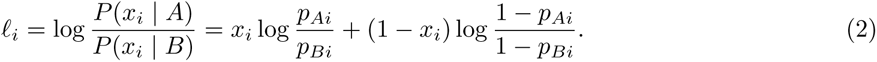

#### Windowed likelihood

The log ratio is summed within non-overlapping windows of 128 segregating sites and the window assigned to the source with the larger total. No information is shared between windows.

#### Likelihood + HMM

The same window scores serve as emissions for a two-state hidden Markov model with a symmetric transition matrix; the switch probability per window is set to *gρs*, where *s* is the mean physical span of a window. Ancestry is decoded by the Viterbi algorithm. This is an RFMix-like baseline in that it applies Markov smoothing to windowed likelihood emissions.

#### Dilated CNN

A residual stack of nine dilated one-dimensional convolution blocks (kernel width 5, dilations 1 through 256, 64 channels, GroupNorm and GELU, 226,177 parameters) maps a four-channel input (target allele, both panel frequencies, and the per-site log likelihood ratio) to a per-site logit. Supplying *ℓ_i_* as an input channel ensures the network holds no information advantage over the likelihood baselines. Training uses binary cross-entropy, AdamW at learning rate 10*^−^*^3^ with weight decay 10*^−^*^4^, cosine annealing, batch size 32, for 15 epochs; held-out accuracy plateaus by epoch 3.

Normalisation is GroupNorm rather than the batch normalisation used in comparable convolutional LAI architectures (Medina Tretmanis et al. 2026). A batch here is a set of haplotypes drawn from the same genomic window, so its members share both the local linkage-disequilibrium background and, through the admixture process, correlated ancestry — the quantity being predicted. Statistics pooled over such a batch are therefore not independent of the label, and normalising by them couples predictions across haplotypes that the task defines as separate instances. Replacing every GroupNorm with batch normalisation, holding the convolution weights and all other hyperparameters fixed, lowers held-out accuracy on CHB/CDX by 0.035 (paired over 5 seeds sharing identical windows, worse on 5 of 5). The coupling is directly visible: scoring the same trained network with normalisation computed from the evaluation batch rather than from running statistics shifts its accuracy by up to 0.008, and the shift changes sign across seeds, whereas the corresponding quantity for GroupNorm is exactly 0.000 because each haplotype is normalised independently. Invariance to batch composition is also a practical requirement here, since the architecture variants we compared differ in memory cost and cannot all be run at one batch size.

### Real-haplotype panels

Source panels were additionally drawn from the 1000 Genomes 30*×* phased release (GRCh38) (Byrska-Bishop et al. 2022), using chromosome 22 between 16 and 51 Mb. Populations are referred to throughout by their 1000 Genomes codes: **CHB** Han Chinese in Beijing; **CHS** Southern Han Chinese; **JPT** Japanese in Tokyo; **CDX** Chinese Dai in Xishuangbanna; **CEU** Utah residents with northern and western European ancestry; **TSI** Tuscans in Italy; **FIN** Finnish in Finland; **GBR** British in England and Scotland; **IBS** Iberian populations in Spain; **GIH** Gujarati Indians in Houston; **PJL** Punjabi in Lahore; **BEB** Bengali in Bangladesh. The eleven pairs span three superpopulations (East Asian, European and South Asian) with six within-superpopulation pairs covering the difficult low-divergence regime and five cross-continental pairs supplying the high-divergence end. Sites were retained if biallelic SNVs, fully called, and polymorphic above 1% minor allele frequency in the pooled sample of both populations; pooling for the frequency filter avoids ascertaining sites on the very frequency difference the inference exploits. Each population contributes 80 reference and 80 disjoint donor haplotypes.

Two departures from the simulation arm follow from working with a single finite dataset. Replication cannot consist of independent draws from a generative process, and instead comes from repeated random reference/donor partitions; this is weaker replication and we report it as such. More importantly, training and evaluation windows are drawn from genomically disjoint segments of the chromosome separated by a buffer, because linkage disequilibrium correlates nearby windows. In the simulation arm each replicate was an independent coalescent realisation, so train/test independence required no such precaution.

This construction is semi-synthetic. CHB and CHS are two sampling locations along a north–south cline rather than the unadmixed sources of a documented admixture event, and the admixture pulse is imposed rather than historical. Real admixed individuals with known local-ancestry ground truth do not exist at this level of divergence, which is the reason simulation is used at all; substituting real haplotypes into an exact-ground-truth construction is the closest available approximation.

### Released implementations

RFMix v2.03-r0 (Maples et al. 2013) and FLARE 0.6.0 (Browning et al. 2023) were run on the identical simulated haplotypes. Haplotypes were paired into pseudo-diploid individuals and written as phased VCFs with a constant-rate genetic map matching the simulated recombination rate; both tools report per-haplotype ancestry, so the pairing is recovered exactly. Reference panels and admixed targets were the same objects supplied to the methods we implemented ourselves.

The two families nevertheless receive those objects in different representations, and we state the asymmetry plainly because it bears on how the comparison should be read. RFMix and FLARE consume the individual reference haplotypes and a genetic map; the three methods we implemented, the network among them, consume panel allele frequencies and no positional information whatsoever, so they treat consecutive segregating sites as equally spaced. The released tools are thus given strictly more, which makes our feasibility conclusions conservative: where every method fails, it is not for want of information supplied to the strongest ones. It also means the comparison is not a controlled test of learned versus classical inference, and we do not present it as one. Both components of the asymmetry are larger on real chromosomes than in simulation: real haplotypes carry more sharing than a clean split generates, and inter-site spacing is more uneven, with a coefficient of variation of 2.16 against 1.03 in simulation and a 99th-to-50th-percentile ratio of 11.9 against 7.1. Either component could therefore contribute to the loss of advantage we report on real data, and our experiments do not separate them.

Two parameter choices were necessary and both favour the released tools. FLARE’s defaults (min-mac=50, min-maf=0.005) discard most sites at our sample size and were relaxed to min-mac=1, min-maf=0. Both tools default to an admixture age far more recent than simulated (8 and 10 generations against a true 30), so each was given the true value; leaving the defaults in place understated both tools’ accuracy. Ancestry-code orientation was verified against the data rather than assumed, since a transposed mapping yields 1 *−* accuracy.

### Haplotype-matching channels

For the channel ablation, four additional inputs were computed per site: the fraction of sites at which the target haplotype agrees with the single best-matching reference haplotype of each population within a *±*64-site window, and the mean of the top five such matches. Raw match fractions concentrate near 0.99 with a standard deviation of about 0.02, an order of magnitude below the scale of the allele and log-ratio channels; left unscaled the network treats them as constant and performs below chance. Each channel is therefore standardised over the window before use.

### Interventions other than input

Before attributing the real-data shortfall to the network’s input we tried to remove it by other means, and Supplemental Table S3 reports each attempt against its own control, paired on seed. A single self-attention layer and a lightweight state-space layer were each inserted after the dilated stack, pooled over the sequence, and compared against the same configuration without it. The discriminant objective adds a linear-discriminant penalty on final-block features to the segmentation loss. Self-supervised pretraining trains the encoder to reconstruct masked contiguous spans of unlabelled real windows before fine-tuning, with a control that receives the same number of *labelled* windows so that any gain cannot be attributed to exposure alone. Simulation pretraining trains on matched-divergence simulated replicates before fine-tuning on the real segment, and is compared against training on the real segment alone. Capacity was varied by width between 32 and 128 channels, a fifteen-fold range in parameter count, in the sweep described in NOTES tuning.md in the repository; those runs are single and select on a pair evaluated elsewhere here, so we treat them as an optimistic bound rather than an estimate and say so in the table.

### Demographic factorial

To identify which simplification in the simulated demography produces the network’s simulated advantage, we crossed two conditions. Continuous symmetric migration at *m* = 5 *×* 10*^−^*^3^ per generation between the two populations after the split replaces the clean split; at that rate the populations sit near migration–drift equilibrium around *F_ST_* = 0.005, which is the structure a cline such as northern/southern Han has and which a recent clean split cannot produce. The empirical recombination map is the PyrhoCHB GRCh38 map for chromosome 22 (Spence and Song 2019), obtained through stdpopsim (Adrion et al. 2020), replacing the uniform rate used elsewhere. Split times differ between conditions (50, 100 and 200 generations without migration, 200, 400 and 800 with it) because migration suppresses *F_ST_* and the migrating arms need longer splits to approach a comparable divergence; they do not reach the top of the range the clean-split arms span, which is why we report a regression controlling for *F_ST_* alongside the condition means. Each of the twelve cells was run under 3 seeds, with the reference panel, training budget and epoch count matched to the simulated sweep of Fig. 2 so that the control arm is comparable with it.

### Viterbi decoding

To separate the network’s representation from its decision rule we decode the same trained logits two ways. The first thresholds each site independently, as reported throughout. The second treats the logits as emissions of a linear-chain model and decodes by Viterbi, with the probability of an ancestry switch between adjacent sites set to *gρd*, where *g* is the generations since admixture, *ρ* the recombination rate and *d* the physical distance between the sites. The transition weights are therefore fixed by the recombination process rather than learned, and no network parameter changes, so any difference is attributable to the decoding alone.

### Implied admixture times

To express tract fragmentation in units a downstream analysis would care about, we convert each method’s tract lengths into the admixture time they imply. For a single pulse *g* generations ago, ancestry tract lengths are approximately exponentially distributed with mean 1*/g* Morgans; with a constant recombination rate, physical length maps directly to genetic length. We therefore report *g*^ = 1*/L̄* over 8 divergence levels. This is the simplest such estimator and real dating methods fit the full length distribution; we quote it to establish the magnitude and direction of the bias, and calibrate it by applying it to the true tracts.

### Phase switch errors

Haplotypes are paired into individuals as they are for VCF export. For each individual, switch positions are drawn from a Poisson process at a specified rate per Mb, and at each switch the two haplotypes are exchanged from that point onward. Ground-truth ancestry labels are exchanged identically, so the labels continue to describe the observed haplotypes; the effect on the task is that observed haplotypes acquire ancestry transitions of non-recombinational origin. Errors are applied independently to both reference panels and to the admixed targets, and to training and test data at the same rate. We verified that per-site allele counts are bitwise identical before and after injection.

### Block-group ablation

To measure the genomic length-scale on which a trained network relies, we exploit the fact that block *i* of the residual stack contributes approximately 4*d_i_* segregating sites of receptive field, so the blocks form a length-scale decomposition. We remove contiguous groups of blocks from the long-dilation end (all blocks with dilation *≥* 8, *≥* 4, and *≥* 2) and re-evaluate without retraining. Since each block enters as *h ← h* + *f* (*h*), removal is exactly the identity and requires no architectural surgery. The decomposition is ordinal rather than metric. GroupNorm computes its statistics over channels and positions within a haplotype, so each block’s output at a given site also depends on summary statistics of the whole window, and the nominal receptive field 4*d_i_* is therefore a lower bound on the length scale a block can access. Removing the long-dilation blocks removes both their convolutional reach and the normalisation statistics computed inside them, so the retention curve reports dependence on those blocks rather than a calibrated measurement of genomic length scale.

Because accuracy is bounded below by chance, raw accuracy drops are not comparable across divergence levels: a model only slightly above chance cannot lose much. We therefore report the fraction of above-chance accuracy retained,

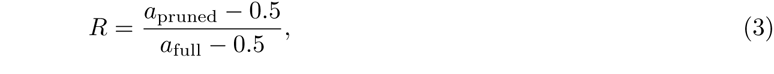

which is interpretable as the share of the model’s usable signal that survives removal of the long-dilation blocks.

This is a measurement of marginal reliance under the trained configuration, not of the minimal sufficient architecture; establishing the latter would require retraining each pruned configuration from initialisation.

### Tract-level metrics

In addition to per-site accuracy we score the shape of the recovered mosaic. For each haplotype we count maximal runs of constant predicted ancestry and compare with the truth, reporting the ratio of tract counts and the ratio of mean tract lengths. We also record, for each true breakpoint, the distance in segregating sites to the nearest predicted breakpoint; as noted in the Results this is only meaningful alongside the tract-count ratio, since over-prediction of breakpoints reduces it artificially. All tract statistics are computed on the same test replicates as the per-site results.

### Evaluation protocol

For each divergence level we simulate 20 training and 8 test replicates with disjoint random seeds. Evaluation windows are 4096 contiguous segregating sites. Training examples are drawn from three random windows per training replicate (3840 haplotype windows); test examples use one deterministic centred window per test replicate (512 haplotype windows). The likelihood baselines are evaluated on the identical windows, so all three methods are scored on the same sites of the same haplotypes. The reported metric is per-site accuracy, the fraction of segregating sites assigned the correct source. For the transfer experiment, each of the 8 trained networks is evaluated against each of the 8 held-out test sets without retraining or recalibration.

### Seed replication

The entire protocol (coalescent simulation, admixture mosaicking, network initialisation and training) is repeated under 3 independent seeds at every divergence level, with simulation seed streams held disjoint across replicates. Error reported in the figures and text is the standard deviation across these independent replicates, which is the quantity that speaks to whether the shape of the accuracy curve is reproducible. Within-run variation across the eight held-out test replicates of a single seed is reported separately for the likelihood baselines and is smaller.

### Implementation and availability

Simulation uses msprime 1.3.4 and tskit 0.6.4; neural network training uses PyTorch 2.8.0 on Apple Silicon via the Metal Performance Shaders backend. Comparisons use RFMix v2.03-r0 and FLARE 0.6.0 as released. Empirical recombination maps are obtained through stdpopsim 0.3.0. Random seeds are fixed and stated in the source, so every figure and every number quoted in the text regenerates exactly from the deposited code.

## Data access

All data underlying this study are publicly available and no new data were generated. Real haplotypes are from the 1000 Genomes Project 30*×* phased release (GRCh38), chromosome 22, obtained from https://ftp.1000genomes.ebi.ac.uk/vol1/ftp/data_collections/1000G_2504_high_coverage/working/20220422_3202_phased_SNV_INDEL_SV/; population assignments are from the Project’s integrated sample panel. Simulated data are not deposited because they are fully determined by the deposited code and the random seeds recorded in it, and regenerate exactly.

All analysis code, the derived result files from which every reported number is generated, and the scripts that build the figures and the manuscript are available at https://github.com/qtianreal/local-ancestry-benchmark under the MIT licence.

## Acknowledgments

This study uses data generated by the 1000 Genomes Project, and we thank the Project and its participants for making the resource publicly available. This research received no specific grant from any funding agency in the public, commercial or not-for-profit sectors.

## Supplemental material

What limits local ancestry inference at low divergence: a feasibility threshold, a metric that conceals failure, and a deficit of input more than architecture

---

**Supplemental Figure S1.**
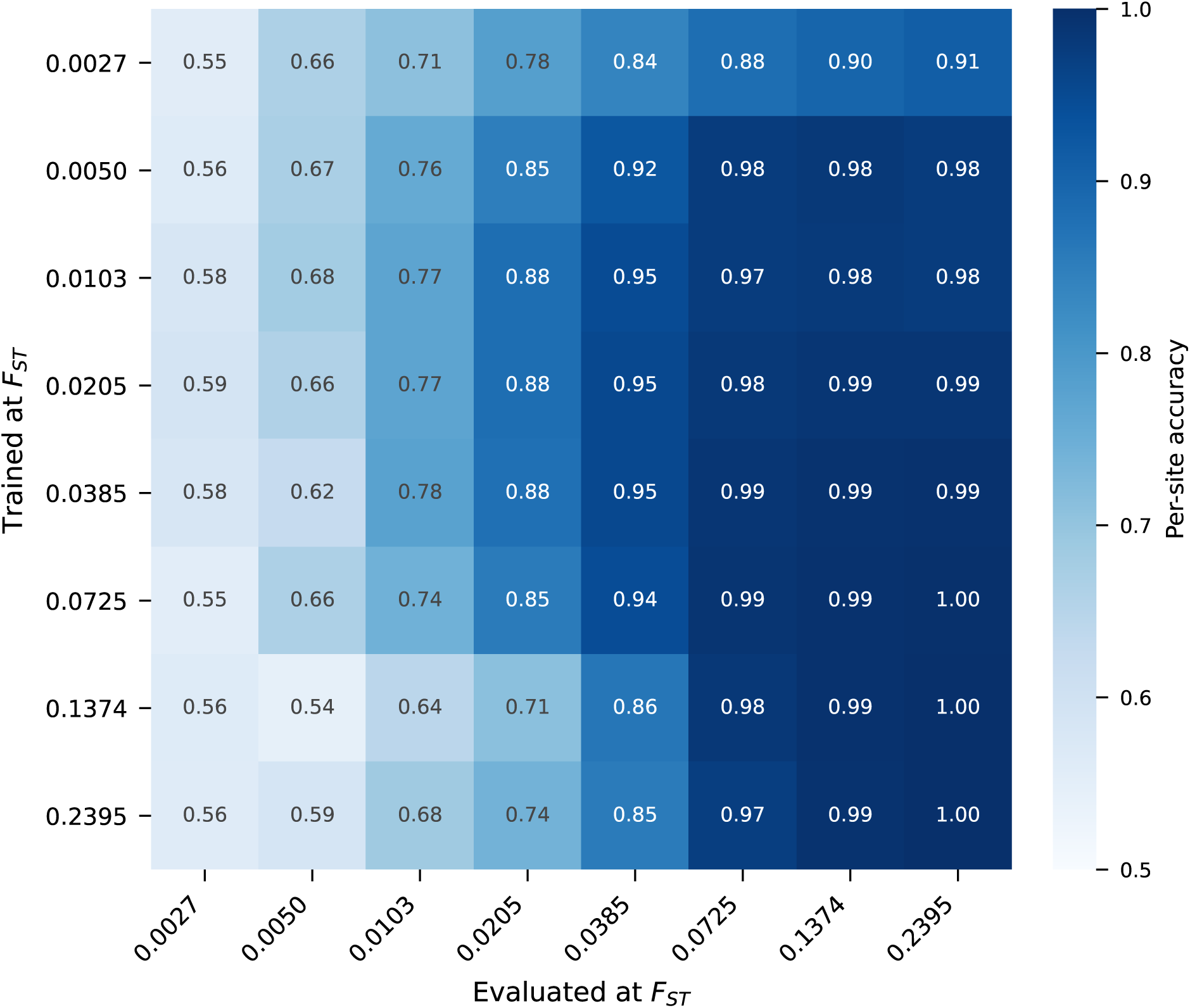
Cross-divergence transfer. Accuracy of each trained network (rows) evaluated on each held-out test set (columns). The diagonal gives matched-divergence performance. Off-diagonal cells quantify degradation under divergence misspecification, the principal robustness concern for simulation-trained methods.

**Supplemental Table S1.**
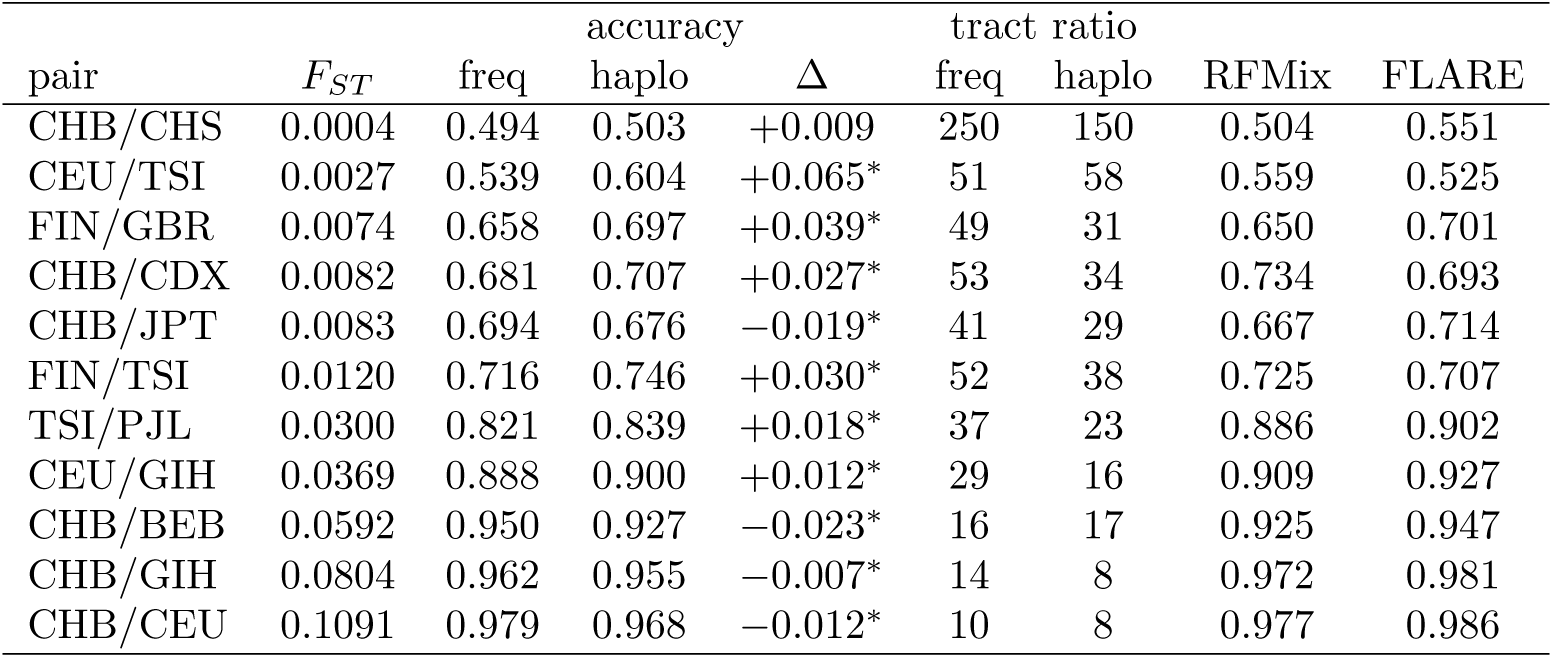
Haplotype features against frequency-only input, on real haplotypes. Per-site accuracy and tract-count ratio (predicted tracts per true tract; 1 is exact), each the mean of five seeds sharing identical partitions, windows and initialisation. Δ is the mean of the paired per-seed differences; *^∗^* marks *p <* 0.05 on a paired *t*-test. RFMix and FLARE are reproduced from the main-text real-data comparison at 80 reference haplotypes and are single runs. Pairs are ordered by measured *F_ST_* ; the sign of Δ changes near *F_ST_* = 0.04.

**Supplemental Table S2.** Accuracy against reference panel size. Per-site accuracy with 40, 80 and 100 reference haplotypes per population. Donors are held at 80 throughout and drawn from the end of the same permutation, so the admixed evaluation data are identical across panel sizes and the reference sets are nested. Network values are means of five seeds; the released tools are single runs. RFMix and FLARE were not run at 40. Every method improves with panel size and none saturates over this range.

| Pair | $F_{ST}$ | Dilated CNN + haplotype | | | RFMix v2 | | FLARE | |
| --- | --- | --- | --- | --- | --- | --- | --- | --- |
|  |  | 40 | 80 | 100 | 80 | 100 | 80 | 100 |
| FIN/GBR | 0.0074 | 0.624 | 0.686 | 0.715 | 0.666 | 0.722 | 0.683 | 0.703 |
| CHB/CDX | 0.0082 | 0.654 | 0.678 | 0.701 | 0.708 | 0.725 | 0.723 | 0.735 |
| CHB/JPT | 0.0083 | 0.676 | 0.718 | 0.738 | 0.770 | 0.786 | 0.766 | 0.775 |
| TSI/PJL | 0.0300 | 0.814 | 0.847 | 0.869 | 0.867 | 0.890 | 0.889 | 0.890 |

**Supplemental Table S3.** Interventions that were not changes of input. Change in per-site accuracy on real haplotypes for each intervention we tried, against its own control, paired on seed. Positive counts are seeds (pairs, for the input row) on which the intervention helped. The controls differ by arm because each is the appropriate one: the attention and state-space arms add a layer to the same trained configuration, the pretraining arms are compared against training on the real segment alone, and the discriminant arm against the same haplotype-aware network without the extra loss term. Capacity is the exception to the pairing: those runs are single, and the sweep they come from selects on a pair evaluated elsewhere in this study, so the value is an optimistic bound rather than an estimate. The final row is the input change, shown on the same scale for comparison. All but the last three rows fall within 0.006 of zero. Of those three, the input change is the one the manuscript recommends; simulation pretraining helps the frequency-only network and not the haplotype-aware one, a contrast interpreted in the Discussion.

| Intervention | Evaluated on | $n$ | $\Delta$ accuracy | Positive |
| --- | --- | --- | --- | --- |
| Self-attention layer | CHB/CDX | 3 | -0.006 | 1/3 |
| State-space layer | CHB/CDX | 3 | +0.006 | 2/3 |
| Capacity, 15-fold parameter range | CHB/CDX | 2 | +0.003 | 2/2 |
| Discriminant objective | TSI/PJL | 5 | -0.001 | 1/5 |
| Self-supervised pretraining | CHB/CDX | 3 | +0.002 | 2/3 |
| More labelled data | CHB/CDX | 3 | -0.004 | 1/3 |
| Simulation pretraining, haplotype-aware | CHB/CDX | 3 | -0.009 | 0/3 |
| Simulation pretraining, frequency-only | CHB/CDX | 3 | +0.061 | 3/3 |
| *Input*: haplotype-matching channels | 8 pairs | 8 | +0.031 | 8/8 |

**Supplemental Table S4.** Demographic factorial. Continuous symmetric migration (*m* = 5 *×* 10*^−^*^3^ per generation) crossed with an empirical recombination map, three split times per condition and three seeds per cell. Split times differ between conditions because migration suppresses *F_ST_* : the migrating arms need longer splits to reach a comparable divergence, and even so they do not span the full range the clean-split arms do, which is why the manuscript quotes a regression controlling for *F_ST_* rather than the raw condition means. Gap is the network’s per-site accuracy minus the better of RFMix and FLARE, computed per run and then averaged. Accuracies are means over the three seeds. The configuration matches the validated simulated sweep (100 reference haplotypes, 12 training replicates, 15 epochs), which an earlier single-seed attempt did not, and which is why that attempt disagreed with the sweep it should have reproduced.

| Condition | $T$ | $F_{ST}$ | Dilated | | FLARE | Gap | SD |
| --- | --- | --- | --- | --- | --- | --- | --- |
|  |  |  | CNN | RFMix v2 |  |  |  |
| Clean split | 50 | 0.0057 | 0.677 | 0.643 | 0.667 | +0.010 | 0.010 |
|  | 100 | 0.0091 | 0.804 | 0.745 | 0.710 | +0.059 | 0.026 |
|  | 200 | 0.0172 | 0.875 | 0.834 | 0.843 | +0.029 | 0.024 |
| + migration | 200 | 0.0057 | 0.602 | 0.603 | 0.615 | −0.026 | 0.030 |
|  | 400 | 0.0050 | 0.635 | 0.657 | 0.574 | −0.022 | 0.027 |
|  | 800 | 0.0050 | 0.615 | 0.603 | 0.570 | +0.010 | 0.060 |
| + empirical map | 50 | 0.0043 | 0.663 | 0.627 | 0.601 | +0.036 | 0.014 |
|  | 100 | 0.0100 | 0.751 | 0.669 | 0.713 | +0.031 | 0.012 |
|  | 200 | 0.0169 | 0.869 | 0.811 | 0.834 | +0.036 | 0.026 |
| + both | 200 | 0.0040 | 0.639 | 0.623 | 0.595 | +0.007 | 0.005 |
|  | 400 | 0.0054 | 0.607 | 0.618 | 0.611 | −0.016 | 0.022 |
|  | 800 | 0.0063 | 0.636 | 0.620 | 0.609 | −0.009 | 0.024 |

